# 2-AG as a potential biomarker for predicting response to short-term fluoxetine treatment in a mouse model of depression

**DOI:** 10.64898/2026.01.14.699270

**Authors:** Andrea Rodríguez-López, El Chérif Ibrahim, Victor Gorgievski, Miquel Martin, Patricia Robledo, Rafael Maldonado, Eleni Tzavara, Beltrán Álvarez-Pérez

**Affiliations:** Laboratory of Neuropharmacology-NeuroPhar, Department of Medicine and Life Sciences, Universitat Pompeu Fabra, Barcelona Biomedical Research Park, 08003 Barcelona, Spain; Laboratory of Neurobiology of Behavior, Department of Medicine and Life Sciences, Universitat Pompeu Fabra, Barcelona Biomedical Research Park, 08003 Barcelona, Spain; Aix-Marseille Univ, CNRS, INT, Inst Neurosci Timone, 13005 Marseille, France; Université Paris Cité, Inserm, CNRS, HealthFex, F-75006 Paris, France; Department of Morphological Sciences, Unit of Human Anatomy and Embryology, Faculty of Medicine, Universitat Autònoma de Barcelona, Bellaterra, Spain; Hospital del Mar Medical Research Institute (IMIM), Barcelona, Spain; Université de Strasbourg (UNISTRA), INSERM UMR-S 1329, Strasbourg Translational Neuroscience and Psychiatry, Centre de Recherche en Biomédecine de Strasbourg, France; Hôpital Sainte Marguerite AP-HM, Pôle de Psychiatrie, 13274 Marseille, France

## Abstract

Major depressive disorder (MDD) is a complex psychiatric condition with significant individual and social impact and marked variability in treatment response. Identifying biomarkers that predict antidepressant efficacy remains a major challenge, and metabolomic approaches offer a powerful tool to uncover biochemical changes associated with depression and treatment outcome. The prefrontal cortex and the hippocampus are key brain regions involved in the pathophysiology of MDD and antidepressant response. Fluoxetine, one of the most commonly prescribed antidepressants, influences not only serotonergic signaling but also the endocannabinoid system. Accordingly, evaluating endocannabinoid levels in these brain regions might provide critical insight into their contribution to mood disorders and the effect of fluoxetine. To identify biomarkers associated with early antidepressant response, male mice were exposed for 5 weeks to the unpredictable chronic mild stress (UCMS) protocol, with fluoxetine treatment administered during the final week. Liquid chromatography–tandem mass spectrometry analyses of hippocampal and prefrontal cortex samples revealed that the depression-like phenotype induced by UCMS was associated with increased hippocampal anandamide (AEA) levels. Behavioral improvement following short-term fluoxetine treatment was accompanied by enhanced hippocampal 2-arachidonoylglycerol (2-AG), which was associated with reduced GABA and glutamate levels in the prefrontal cortex. These findings suggest that increased 2-AG levels may serve as a predictor of early response to fluoxetine.

## Introduction

Depressive disorders include a cluster of diseases characterized by the presence of sad, empty, and/or irritable mood, associated with alterations that significantly affect the individual’s ability to function ^1^. Endocannabinoids are central in modulating various neurotransmitter systems ^2^, including glutamatergic ^3^, GABAergic, and monoaminergic ^4,5^ signaling, all of which are implicated in the pathogenesis of major depressive disorder (MDD) ^6^. Altered levels of the arachidonic acid derivatives, anandamide (AEA) and 2-arachidonoylglycerol (2-AG), have been associated with stress vulnerability and impaired emotional regulation ^2,7^. According to its duration and effects on daily activity, MDD is the most disruptive mental health problem ^8^. Almost half of the patients do not respond to antidepressant medication ^9,10^, which is associated with increased healthcare utilization and cost ^11^. Treatments avoiding relapse and achieving full remission remain a major challenge. Selective serotonin reuptake inhibitors (SSRIs), including fluoxetine, are considered a first-line option ^12^. Fluoxetine has high acceptability ^13,14^ and beneficial off-target effects on noradrenaline and dopamine ^15^, inflammation ^16^ and the endocannabinoid system ^17,18^. However, it has a high variability of response ^13^, which is a setback in clinical practices, where every additional treatment step decreases the probability of remission and response success ^19^. Unlike other SSRIs, fluoxetine also increases noradrenaline and dopamine extracellular levels in the prefrontal cortex ^15^ through the blockade of 5-HT2C receptors on cortical interneurons ^20^. Nonetheless, it is crucial to differentiate acute and chronic effects. Acute fluoxetine increases extracellular serotonin by blocking the transporter ^15,21,22^. However, repeated treatment causes an excess of extracellular serotonin that triggers 5-HT1A autoreceptor activity, which subsequently reduces serotonin synthesis and turnover ^22,23^ and lower serotonin intracellular levels ^21^. Yet, chronic fluoxetine treatment (> 2 weeks) leads to 5-HT1A autoreceptor desensitization ^23^ and restoration of serotonin intracellular levels ^21^. Advanced metabolomic techniques, such as liquid chromatography-tandem mass spectrometry (LC-MS/MS), would provide new insight into the role of serotonin and other neurotransmitters in mood disorders and rapid molecular changes in response to fluoxetine. Therefore, the identification of biomarkers of early response to fluoxetine could potentially facilitate and support clinical practice in psychiatry. The unpredictable chronic mild stress (UCMS) model is a reliable mouse model that closely mimics the behavioral deterioration and molecular alterations observed in MDD patients ^24^. Our objective was to identify metabolomic alterations associated with short-term fluoxetine response, in two key brain areas playing a key role in this disorder, the hippocampus and the prefrontal cortex ^25–28^.

## Material and methods

### Animals and ethics

Male C57BL/6J (8 weeks old) (n = 60) were purchased (Charles River, France). Except for control (n = 8) (CONTROL-VEH), which were grouped in 4, mice were individually housed in Plexiglas cages and maintained in an environment with controlled temperature (21L±L1L°C) and humidity (55L±L10%). Food and water were available *ad libitum*. All behavioral experiments were performed during the light phase of a 12-hour cycle (lights on at 7.30 am and off at 7.30 pm). Animal procedures and care were conducted under the European Communities Directive 2010/63/EU and the Spanish Royal Decree 53/201RD, following “Animals in Research: Reporting Experiments” (ARRIVE) guidelines, and approved by the local (Comitè Ètic d’Experimentació Animal-Parc de Recerca Biomèdica de Barcelona, CEEA-PRBB) and autonomous (Generalitat de Catalunya, Departament de Territori i Sostenibilitat) ethical committee. All behavioral and biochemical experiments were performed by a researcher blind to experimental conditions.

### Drug administration

Fluoxetine hydrochloride (0927, Tocris) dissolved in saline (0.9 % NaCl) or vehicle (0.9 % NaCl) was injected intraperitoneally (i.p.) at 10 mg/kg daily from 2 pm to 4 pm for 7 consecutive days.

### Unpredictable chronic mild stress protocol

After a week of acclimation, mice were subjected to 5 weeks of UCMS as previously described^29^. Stressors were applied twice daily during a 3-h period and overnight in a randomized order. Daytime stressors included: wet bedding, 45°-tilted cages, pair or crowded housing, water deprivation, and confinement in a reduced space with a grid surface, restricted food access, and cage interchange. Overnight stressors included: flashing lights, inversion of the light-dark cycle, 45°-tilted cages, and wet bedding. Before the start of the protocol and once a week afterwards, the animals’ weight and apathy (self-grooming and coat state) were evaluated. After 4 weeks, animals under the UCMS protocol were randomly assigned to fluoxetine (UCMS-FLUOX, n = 40) or vehicle (UCMS-VEH, n = 10) groups (Figure 1-(a)). At the end of the experimental protocol and 24 h after the last administration, animals were euthanized by cervical dislocation. After decapitation, brains were removed, placed over ice, and the prefrontal cortex and hippocampus were dissected, weighed, frozen in liquid nitrogen, and stored at -80°C.

**Figure 1.**
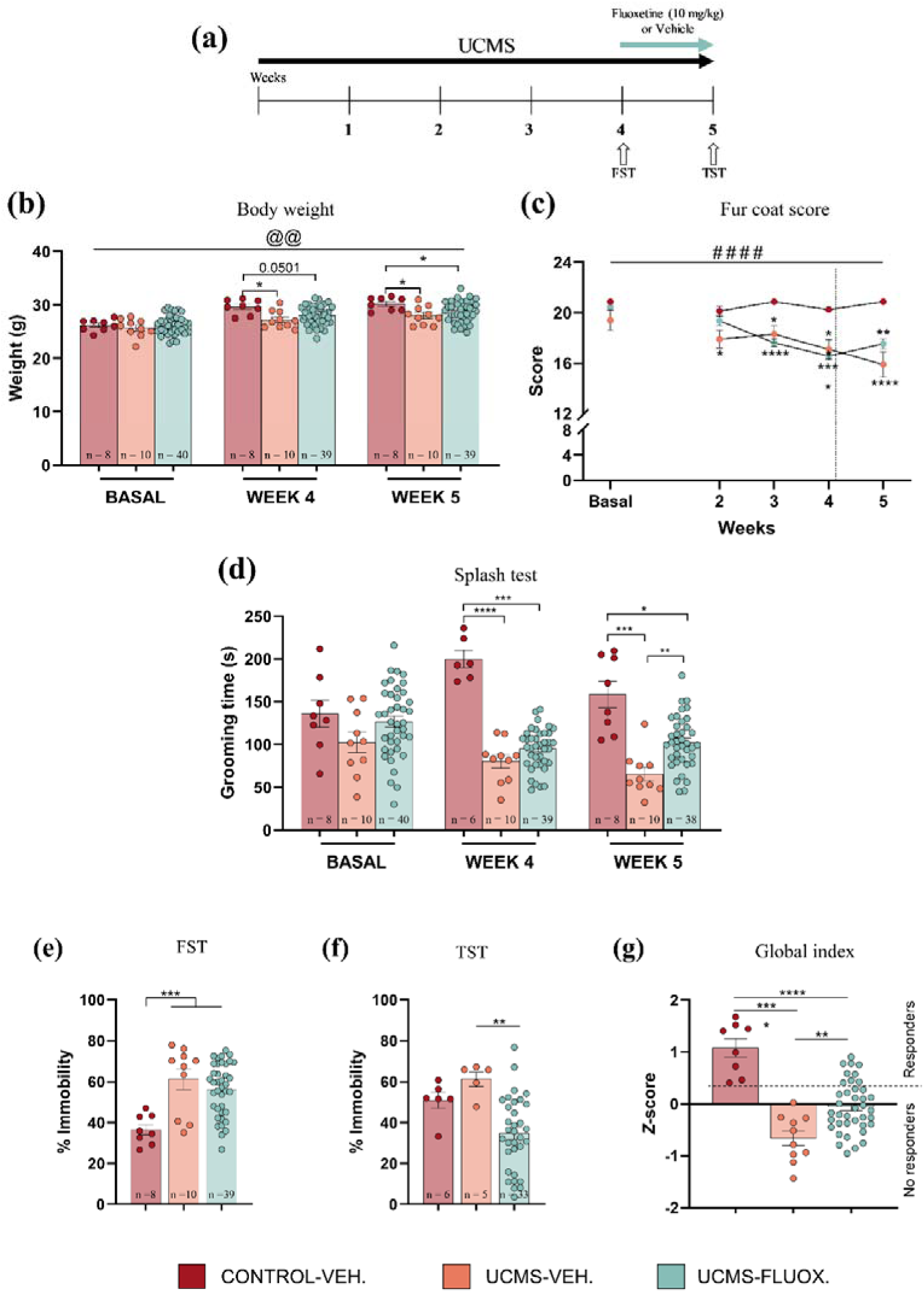
Effects of short-term fluoxetine treatment on depressive-like behavior. **(a)** Schematics of the UCMS protocol. **(b)** Body weight measures, **(c)** fur coat score, and **(d)** grooming time in the splash test, during UCMS and after 1 week of treatment with fluoxetine. Depressive-like behavior **(e)** before starting fluoxetine treatment in the FST and **(f)** after 7 days of treatment in the TST. **(g)** Computation of a global index from the z-score value of total time grooming, body weight, and fur coat score at the end of the experimental protocol. Data are expressed as mean ± s.e.m. The vertical dashed line in panel (c) indicates the start of fluoxetine or vehicle treatment. Individual data points are depicted as circles, and n = indicates the number of male mice per group. The dashed vertical line in (c) indicates the starting point of fluoxetine treatment. * p < 0.05, ** p < 0.01, **** p < 0.001; #### p < 0.001 (time); @@ p < 0.01 (time x group) by one-way ANOVA or mixed-effect analysis followed by Bonferroni *post-hoc*.

### Apathy

Fur coat and grooming time were measured as indicators of apathy ^24^. The score of the animal’s coat state was calculated as the sum of scores, one for each of 7 different body parts (head, neck, dorsal, ventral, tail, forepaws, hind paws). Each part was given a score on a 3-point scale (1 = poor, 2 = fair, 3 = very good) (26). In the splash test, nesting material was temporarily removed, and animals were splashed with a 20 % glycerol solution on the dorsal surface. Self-directed grooming behavior was recorded for 5 min, and latency, total number of events, and time spent self-grooming were manually measured.

### Behavioral despair

Behavioral despair was measured with the forced swim test (FST) ^30^ after 4 weeks of UCMS and the tail suspension test (TST) at the end of the experimental protocol. In the FST, mice were placed in water-filled cylinders (30 cm high and 15 cm diameter) for 6 min. Immobility time, considered when mice floated in an upright position and made only small movements to keep their head above, was manually measured during the last 4 min ^31,32^. In the TST, mice were individually suspended by the tail using adhesive tape placed 1 cm from the tip of the tail and recorded during a 6-min session. Immobility time, considered as the complete absence of movement, was registered from the last 4 min ^31^. Animals falling or climbing over their tails were discarded.

### Quantification of brain endocannabinoids by LC-MS/MS

AEA and 2-AG were quantified as previously described ^33^ with modifications. Briefly, one hippocampal hemisphere (21.2 ± 6.43 mg) was homogenized in 700 µL of 50 mM Tris–HCl (pH 7.4):methanol (1:1) containing deuterated internal standards (AEA-d4, DHEA-d4, 2-AG-d5, and 2-OG-d5; Cayman Chemical and Toronto Research Chemicals). Chloroform (5 mL) was added, samples were shaken for 20 min and centrifuged (1700 g, 5 min, room temperature). The organic phase was evaporated under nitrogen, reconstituted in 100 µL water:acetonitrile (10:90, v/v) with 0.1% formic acid, and transferred to microvials for LC-MS/MS analysis.

Analyses were performed using a Waters Acquity UPLC system coupled to a Xevo TQ-S micro mass spectrometer. Chromatographic separation was achieved on a BEH C18 column (2.1 × 100 mm, 1.8 µm) at 40°C, with a flow rate of 0.4 mL/min and a mobile phase consisting of 0.01% formic acid in water and acetonitrile. Detection was carried out in multiple reaction monitoring (MRM) mode, and quantification was performed by isotope dilution.

### Quantification of brain metabolites by LC-MS/MS

The remaining hippocampal tissue (21.06 ± 7.16 mg) and prefrontal cortex samples (26.35 ± 8.99 mg) were homogenized in acetonitrile:water (2:1, 0.1% formic acid; 25 mg tissue/600 µL). After centrifugation (14,500 rpm, 5 min, 4°C), supernatants were collected and processed using two analytical procedures.

For GABA and glutamate determination (method A), aliquots (10 µL) were mixed with labelled internal standards (GABA-d6 and Glu-d5) and acetonitrile, vortexed, centrifuged, and transferred to microvials. For monoamines, amino acids, and metabolites (method B), larger aliquots (300 µL) were combined with a labelled internal standard mixture, evaporated under nitrogen, and derivatized with dansyl chloride in alkaline conditions prior to LC-MS/MS analysis.

Analyses were conducted on an Acquity I-Class UPLC coupled to a Xevo TQ-S micro mass spectrometer equipped with an electrospray ionization source operating in positive mode. Source parameters were set as follows: capillary voltage 3 kV, source temperature 150°C, desolvation temperature 600°C, desolvation gas flow 1200 L/h, and cone gas flow 50 L/h. Nitrogen was used as nebulizing and drying gas, and argon as collision gas. Data acquisition and processing were performed using MassLynx v4.1.

For method A, separation was achieved on a BEH Amide column using an acetonitrile/water gradient containing ammonium formate and formic acid. For method B, a BEH C18 column and a methanol/water gradient were employed. MRM transitions for each analyte are reported in Table 1. Calibration curves were generated at 12 concentration levels using external standards and processed with TargetLynx XS software.

**Table 1.** MRM transitions used for method B.

| Analyte | Precursor ion m/z | CV (V) | Product ion m/z | CE (eV) |
| --- | --- | --- | --- | --- |
| 5-HT | 643 | 15 | 170, 408 | 35, 30 |
| DA | 853 | 15 | 170, 619 | 45, 35 |
| NA | 869 | 15 | 171, 617 | 45, 35 |
| Gly | 309 | 15 | 157, 170 | 30, 15 |
| 5-HIAA | 425 | 15 | 171, 381 | 30, 20 |
| DOPAC | 635 | 15 | 170, 401 | 30, 30 |
| HVA | 416 | 15 | 156, 170 | 30, 20 |
| 3-MeO-Tyr | 634 | 15 | 170, 400 | 35, 35 |
| Phe | 399 | 15 | 120, 157 | 20, 30 |
| Tyr | 648 | 15 | 369, 414 | 30, 30 |
| Trp | 438 | 15 | 159, 171 | 30, 30 |
| 5-HT-d4 | 647 | 15 | 412 | 30 |
| DA-d4 | 857 | 15 | 623 | 35 |
| NA-d6 | 875 | 15 | 623 | 35 |
| Gly-13C2,15N | 312 | 15 | 157 | 30 |
| 5-HIAA-d4 | 429 | 15 | 171 | 30 |
| DOPAC-d3 | 638 | 15 | 170 | 30 |
| HVA-13C,d3 | 420 | 15 | 156 | 30 |
| 3-MeO-Tyr-d4 | 638 | 15 | 170 | 35 |
| Phe-d5 | 404 | 15 | 157 | 30 |
| Tyr-d4 | 652 | 15 | 373 | 30 |
| Trp-d5 | 443 | 15 | 159 | 30 |

### Statistical analysis

Behavioral and metabolomic results are presented as mean ± standard error of the mean (s.e.m). Based on previous evidence that distinguishes fluoxetine treatment responders from non-responders ^14,34^, a global index score was calculated using Z-normalized values of total grooming time, fur coat score, and body weight. The UCMS-FLUOX group was classified into responders and non-responders according to the lowest global index value observed in CONTROL-VEH mice. Serotonin and dopamine turnover were estimated using their respective metabolite ratios. Data normality was assessed using the Shapiro–Wilk test. Statistical analyses were performed using two-way ANOVA or mixed-effects models for repeated measures, with significance set at *p* < 0.05. When appropriate, Bonferroni or Fisher’s LSD post hoc tests were applied. Outliers, defined as values exceeding ±2 standard deviations from the mean, were excluded. Data analysis and visualization were conducted using IBM SPSS v23 and GraphPad Prism v10. Pearson’s correlation analyses were performed in R (v4.4.1) and visualized using the ggcorrplot package (v0.1.4.1).

## Results

### Effects of short-term exposure to fluoxetine on stress-induced depressive-like behavior

Physical condition and behavior were evaluated to assess the effects of UCMS and short-term fluoxetine treatment. For body weight, time, and group differences after mixed-effects analysis were observed (time x group, F[4, 108] = 4.287, *p* = 0.0029). Animals exposed to UCMS showed significantly lower body weights than control mice (CONTROL-VEH. *vs.* UCMS-VEH. *p* = 0.0123) after 4 weeks. Fluoxetine did not significantly reverse these effects (CONTROL-VEH. *vs.* UCMS-VEH. *p* = 0.0181; CONTROL-VEH. *vs.* UCMS-FLUOX. *p* = 0.0383) (Figure 1-(b)). There were fur coat time and group differences (time: F[3.333, 181.6] = 15.17, *p* < 0.0001; group: F[2, 55] = 9.347, *p* = 0.0003; time x group: F[8, 218] = 6.133, *p* < 0.0001). Fur was significantly degraded in stressed animals after 4 weeks (CONTROL-VEH. *vs.* UCMS-VEH. *p* = 0.0086; CONTROL-VEH. *vs.* UCMS-FLUOX. *p* < 0.0001), and fluoxetine had no influence (Figure 1-(c)). Mixed-effects analysis showed significant group differences in the total grooming time (F[2, 55] = 32.61, p < 0.0001). Bonferroni’s multiple comparison test exposed no basal differences, but statistically significant differences before treatment between stressed and control animals (CONTROL-VEH. *vs.* UCMS-VEH. *p* < 0.0001, CONTROL-VEH. *vs.* UCMS-FLUOX. *p* = 0.0001). Fluoxetine partially increased total grooming time (UCMS-VEH. *vs.* UCMS-FLUOX. *p* = 0.0033), although not comparable to control animals (CONTROL-VEH. *vs.* UCMS-FLUOX. *p* = 0.0252) (Figure 1-(d)). Immobility time was increased in UCMS-exposed animals (one-way ANOVA F[2, 54] = 9.42, *p* = 0.0003; CONTROL-VEH.: *vs.* UCMS-VEH. *p* = 0.0005; and *vs.* UCMS-FLUOX. *p* = 0.0007) (Figure 1-(e)). Short-term fluoxetine treatment significantly decreased immobility time in the TST (one-way ANOVA F[2, 41] = 7.153, *p* = 0.0022, UCMS-VEH. *vs.* UCMS-FLUOX. *p* = 0.0052) (Figure 1-(f)). One-way ANOVA in global index (Figure 1-(g)) revealed significant group differences (F[2, 54] = 29.14, *p* < 0.0001; CONTROL-VEH. *vs.* UCMS-VEH. *p* < 0.0001, CONTROL-VEH. *vs.* UCMS-FLUOX. *p* < 0.0001; UCMS-VEH. *vs.* UCMS-FLUOX. *p* = 0.0027). Therefore, UCMS induced significant physical and emotional deterioration, which was only partially reversed by short-term fluoxetine treatment. The global index helped distinguish responders from non-responders to treatment.

### UCMS exposure and short-term fluoxetine modulate endocannabinoid levels in the hippocampus

Stress and fluoxetine treatment significantly influenced endocannabinoids in the hippocampus (one-way ANOVA for 2-AG: F[3, 51] = 5.645, *p* = 0.002; and AEA: F[3,51] = 6.582, *p* = 0.0008) (Figure 2-(a), and (b)). Behavioral improvement by short-term fluoxetine treatment in responders was associated with increased 2-AG levels (*p* = 0.0016 to CONTROL-VEH.; *p* = 0.0005 to UCMS-VEH.; *p* = 0.0123 to non-responders) (Figure 2-(a)). In UCMS-VEH. animals, AEA levels were significantly increased by stress alone (*p* = 0.0198 to CONTROL-VEH.), whereas fluoxetine decreased them in both responders and non-responders (responders: *p* < 0.0001; and non-responders *p* = 0.0007) (Figure 2-(b)).

**Figure 2.**
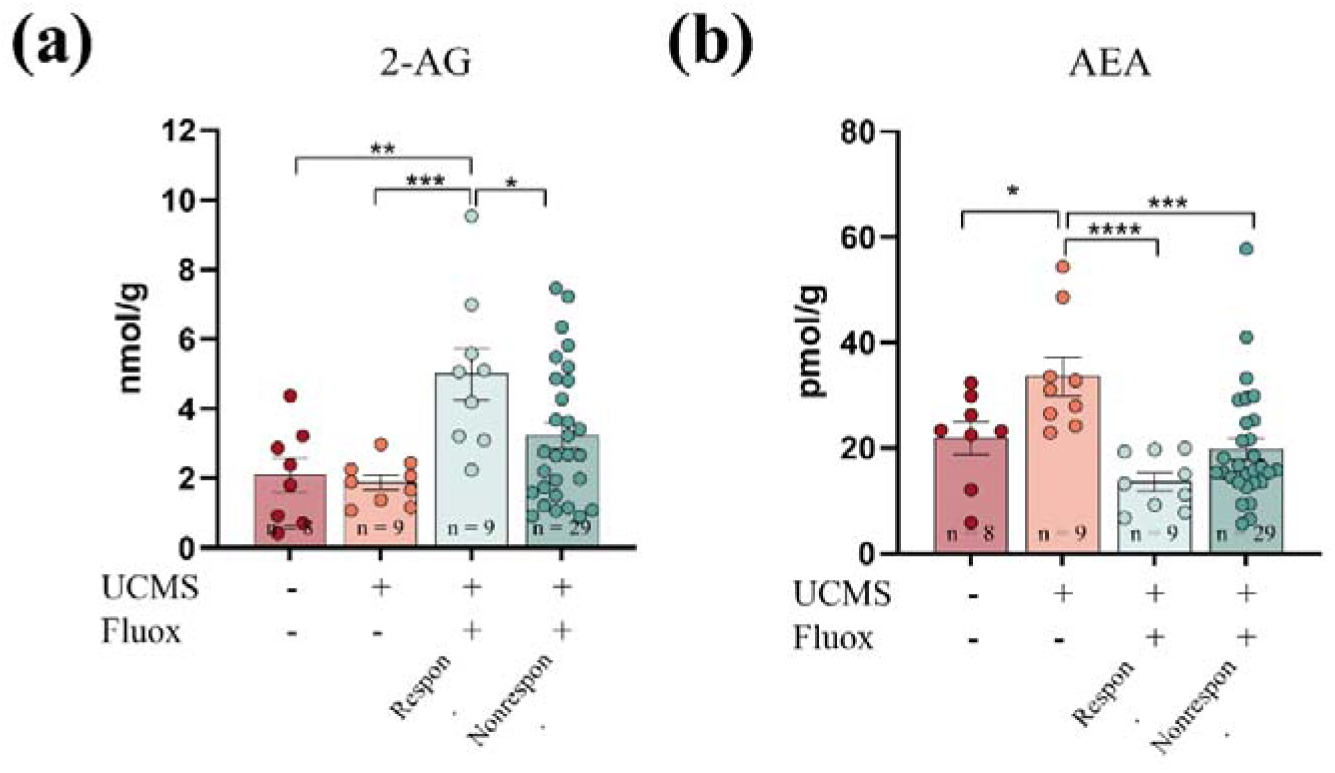
Effects of UCMS and short-term treatment with fluoxetine on hippocampal levels of endocannabinoids. Endocannabinoids **(a)** 2-AG and **(b)** AEA quantified by LC-MS/MS. UCMS-subjected mice treated with fluoxetine are separated into responders (Respon.) and non-responders (Nonrespon.). Data are expressed as mean ± s.e.m. Individual data points are depicted as circles and n = indicates the number of mice per group. * p < 0.05, ** p < 0.01, *** p < 0.001, **** p < 0.0001; by one-way ANOVA followed by uncorrected LSD *post-hoc*.

### UCMS and short-term fluoxetine exposure modulate neurotransmitter levels in the hippocampus and the prefrontal cortex

Fluoxetine treatment significantly decreased the serotonin pool (one-way ANOVA F[3, 51] = 4.456, *p* = 0.0074) (Figure 3-(a)) and turnover (one-way ANOVA F[3, 50] = 27.29, *p* < 0.0001) (Figure 3-(b)). Dopamine also decreased in responders compared to CONTROL-VEH. animals (one-way ANOVA F[3, 51] = 3.725, *p* = 0.0169, *p* = 0.0221) (Figure 3-(c)), but the turnover was not affected (Figure 3-(d)). There were no differences in noradrenaline (Figure 3-(e)), glutamate (Figure 3-(f)), GABA (Figure 3-(g)), nor glycine (Figure 3-(h)) in the hippocampus. Stress does not influence neurotransmitters in the hippocampus.

**Figure 3.**
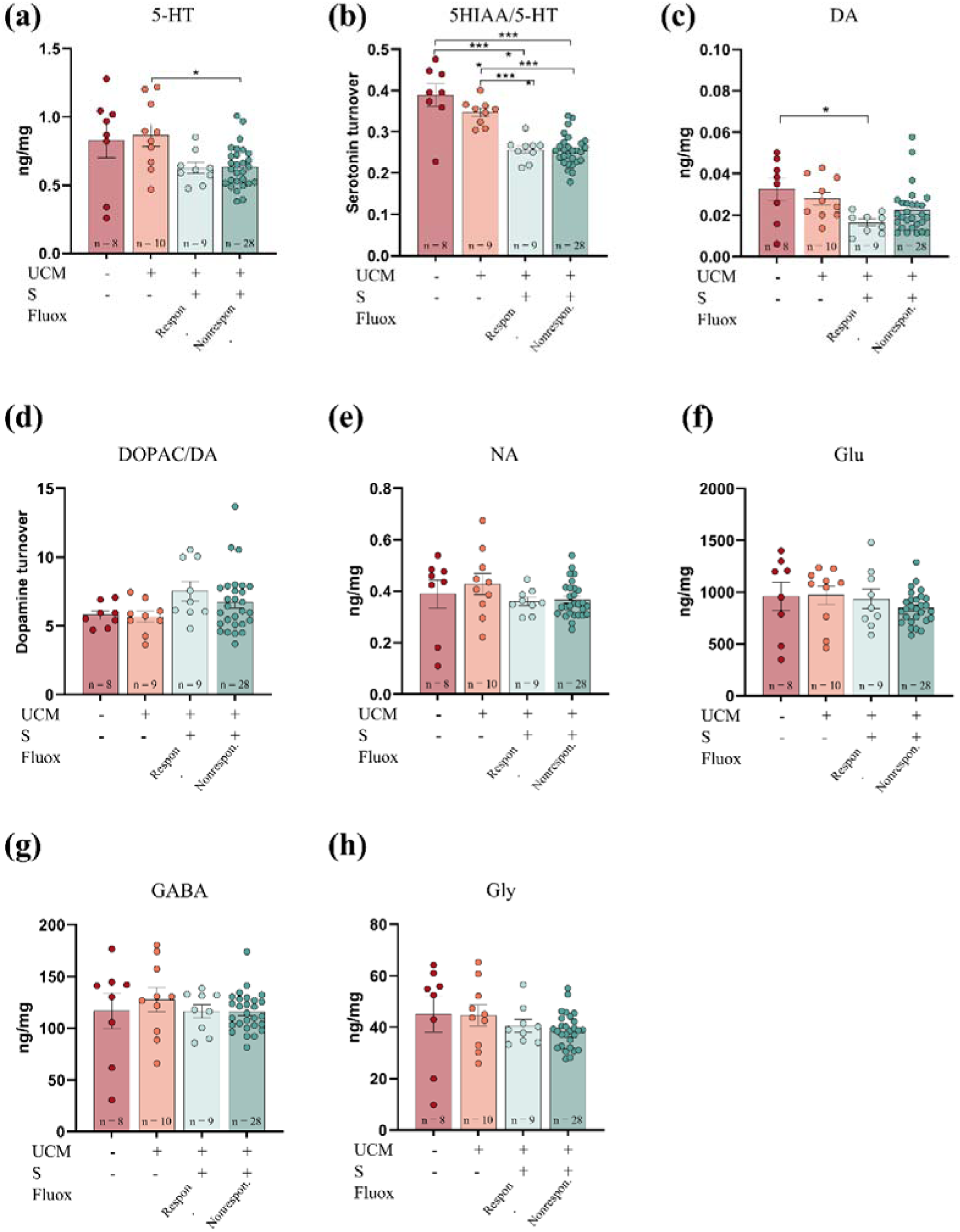
Effects of UCMS and short-term fluoxetine treatment on hippocampal levels of neurotransmitters and related metabolites. **(a**) Serotonin (5-HT) and **(b)** serotonin turnover (5-HIAA/5-HT), **(c**) dopamine (DA) and **(d)** dopamine turnover (DOPAC/DA), **(e)** noradrenaline, **(f)** glutamate (Glu), **(g)** GABA and **(h)** glycine quantification by LC-MS/MS. Stressed mice treated with fluoxetine were separated into responders (Respon.) and non-responders (Nonrespon.). Data are expressed as mean ± s.e.m. Individual data points are depicted as circles and n = indicates the number of male mice per group. * p < 0.05, ** p < 0.01, *** p < 0.001, **** p < 0.0001; by one-way ANOVA followed by Bonferroni *post-hoc*.

Stress did not significantly alter serotonin nor the turnover in the prefrontal cortex, while fluoxetine significantly decreased them (levels: one-way ANOVA F[3, 50] = 10.47, responders and non-responders to CONTROL-VEH. *p* < 0.0001, and to UCMS-VEH. *p* < 0.0001; turnover: one-way ANOVA F[3, 50] = 42.83, *p* < 0.0001, to CONTROL-VEH. responders *p* = 0.0233, and non-responders *p* = 0.0172, to UCMS-VEH. responders *p* = 0.0233, and non-responders *p* = 0.0172) (Figure 4-(a) and (b)). Oppositely, dopamine and its turnover were not altered by stress or by short-term fluoxetine (Figure 4-(c) and (d)). As seen in the hippocampus, noradrenaline levels are unmodified by chronic stress or fluoxetine (Figure 4-(e)). In the prefrontal cortex, glutamate was decreased by fluoxetine in responders (one-way ANOVA F[3, 50] = 4.231, *p* = 0.0097, *vs.* CONTROL-VEH.: *p* = 0.0094, *vs.* UCMS-VEH.: *p* = 0.0019, *vs.* non-responders *p* = 0.0476 by uncorrected Fisher’s LSD) (Figure 4-(f)). Likewise, GABA was decreased by fluoxetine in responders (one-way ANOVA F[3, 50] = 4.489, *p* = 0.0072, vs. CONTROL-VEH *p* = 0.0233; vs. UCMS-VEH *p* = 0.0009), with a marked tendency compared to non-responders (*p* = 0.0594) (Figure 4-(g)). On the other hand, glycine levels were significantly decreased by fluoxetine to the same extent in responders and non-responders (one-way ANOVA F[3, 50] = 4.633, *p* = 0.0062; UCMS-VEH. vs. responders *p* = 0.0198; vs. non-responders *p* = 0.006) (Figure 4-(h)).

**Figure 4.**
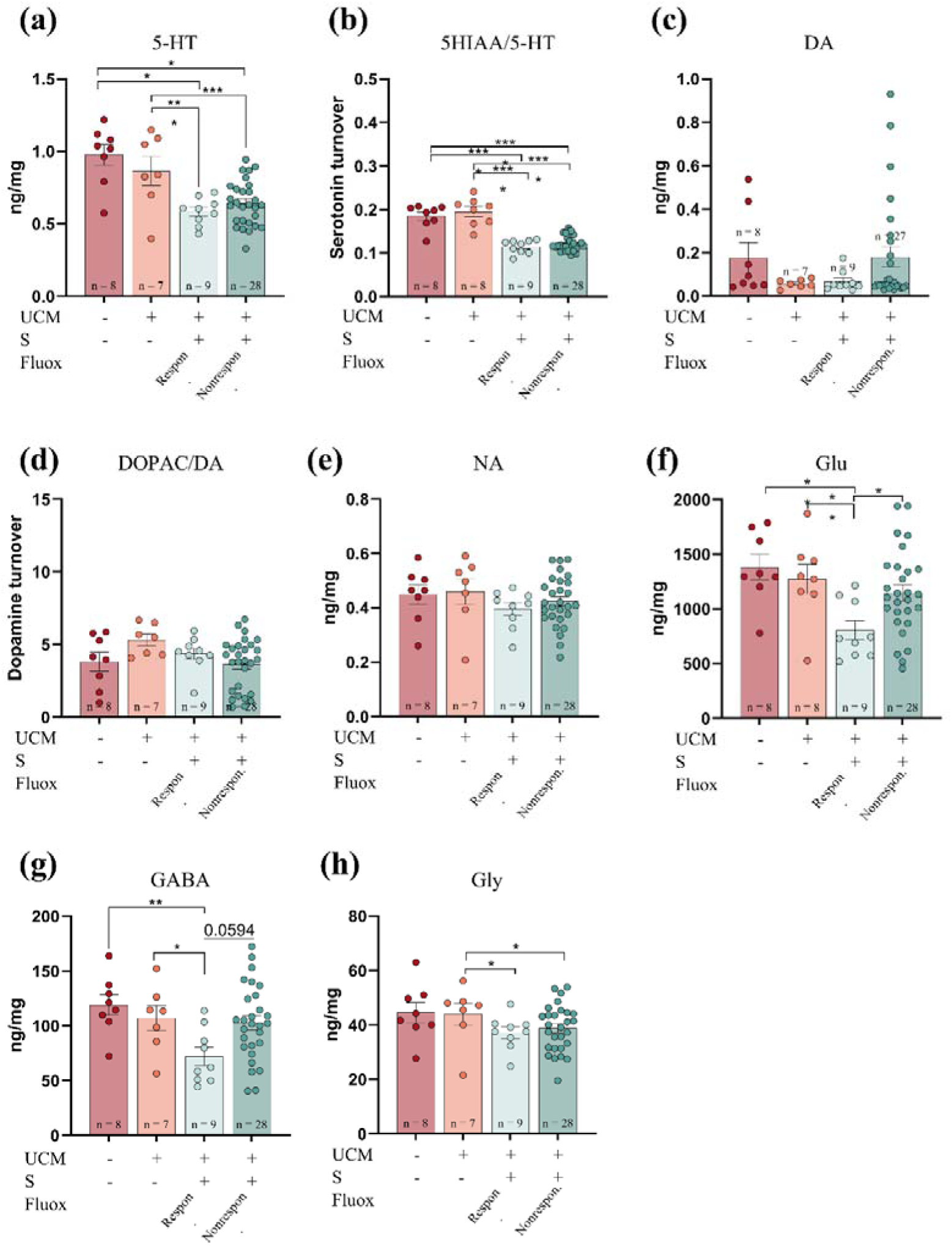
Effects of UCMS and short-term fluoxetine treatment on prefrontal cortex levels of neurotransmitters and related metabolites. **(a**) Serotonin (5-HT) and **(b)** serotonin turnover (5-HIAA/5-HT), **(c**) dopamine (DA) and **(d)** dopamine turnover (DOPAC/DA), **(e)** noradrenaline, **(f)** glutamate (Glu), **(g)** GABA and **(h)** glycine quantification by LC-MS/MS. Stressed mice treated with fluoxetine are separated into responders (Respon.) and non-responders (Nonrespon.). Data are expressed as mean ± s.e.m. Individual data points are depicted as circles, and n = indicates the number of mice per group. * p < 0.05, ** p < 0.01, *** p < 0.001, **** p < 0.0001; by one-way ANOVA followed by Uncorrected LSD (panels f and g) and Bonferroni (panels a-e, and h) *post-hoc*.

### Correlations between neurotransmitter levels in the hippocampus and prefrontal cortex

Neurotransmitter interactions are altered in stress and after antidepressant treatment. We analyzed how UCMS and fluoxetine affect them in the hippocampus and prefrontal cortex.

Pearson correlations revealed that under control conditions, the concentrations of most neurotransmitters in the hippocampus were significantly positively correlated among them (Figure 5-(a)) and were significantly reduced in stressed mice (Figure 5-(b)). Thus, endocannabinoid concentrations no longer correlated with monoamines or excitatory and inhibitory neurotransmitters; the only remaining correlations were between serotonin and monoamines, GABA, glutamate, and glycine. Fewer associations between neurotransmitter levels were observed in the prefrontal cortex of control mice (Figure 5-(a)), while the number of correlations was higher in stressed mice, mostly between inhibitory and excitatory neurotransmitters and with serotonin (Figure 5-(b)).

**Figure 5.**
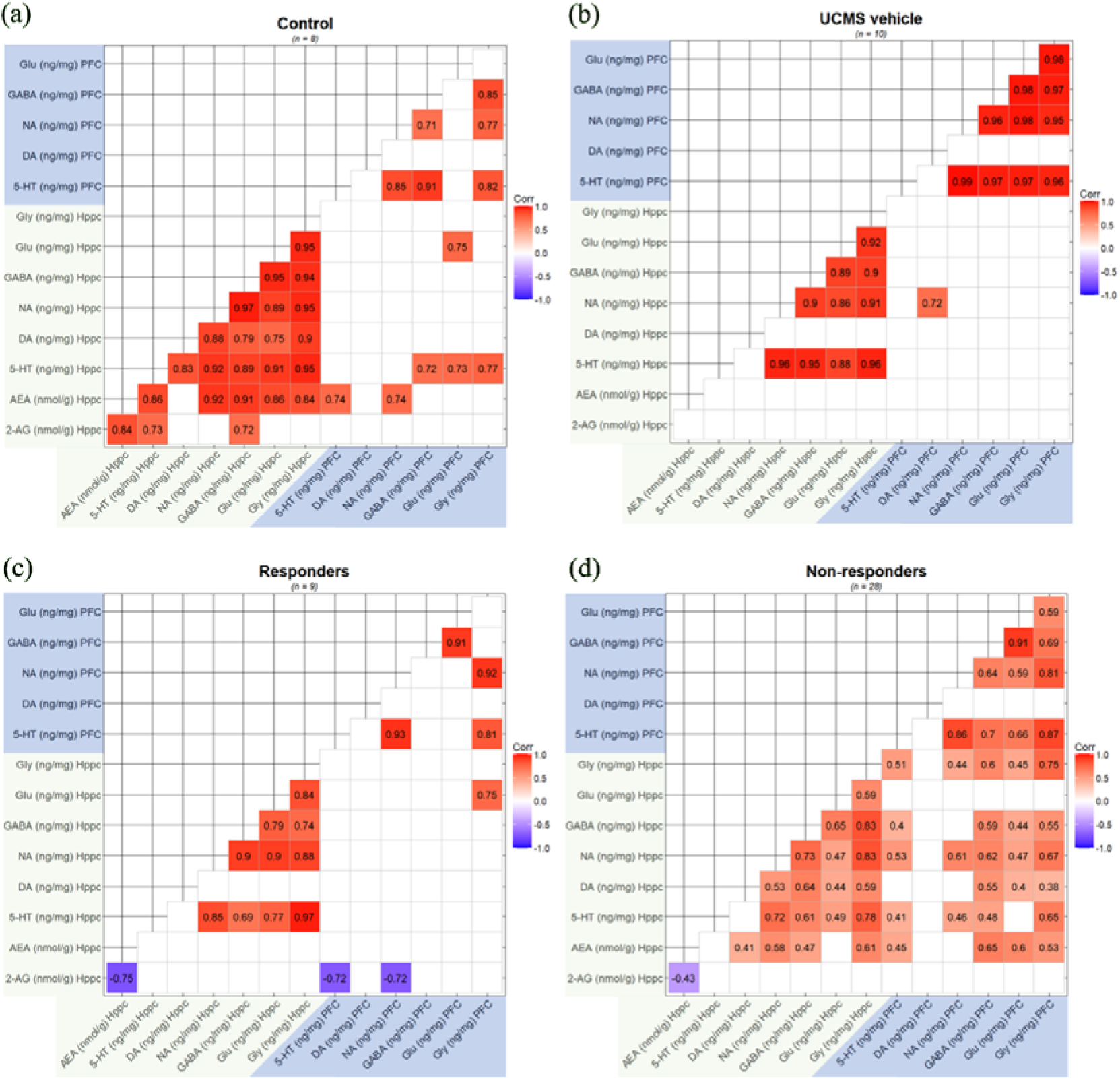
Correlation matrices of neurotransmitters in hippocampus and prefrontal cortex. Each panel illustrates the significant (p<0.05) correlations found in the hippocampus and the prefrontal cortex between all the neurotransmitter concentrations studied. **(a**) control mice, **(b**) stressed mice treated with vehicle, **(c**) responders to fluoxetine, and **(d)** non-responders to fluoxetine. The numbers inside the squares represent the Pearson correlation coefficient (r). The colors inside the squares indicate the strength and direction of the association, with red and blue for positive and negative correlations, respectively.

Short-term fluoxetine induced a distinct pattern of associations in the hippocampus of responders and non-responders. In responders, strong positive correlations were observed, mainly in the hippocampus. Notably, 2-AG concentrations correlated negatively with AEA in the hippocampus and with serotonin and noradrenaline in the prefrontal cortex (Figure 5-(c)). In contrast, in non-responders, dopamine positively correlated with all the other neurotransmitters in the hippocampus (except serotonin) but not in the prefrontal cortex (Figure 5-(d)). In addition, the strong negative correlation between 2-AG and AEA found in the hippocampus of responders is diminished, and the negative correlations between 2-AG and serotonin and noradrenaline in the prefrontal cortex are lost. In contrast to what was observed in responders, AEA concentrations positively correlate with most of the hippocampal prefrontal cortex neurotransmitters (Figure 5-(d)). In the prefrontal cortex of responders, a few significant associations between neurotransmitters were observed (Figure 5-(c)), opposite to non-responders, where most neurotransmitters were correlated with each other in the prefrontal cortex and with neurotransmitters in the hippocampus (Figure 5-(d)).

Stress disrupted hippocampal neurotransmitter associations while increasing correlations in the prefrontal cortex. Fluoxetine partially restored hippocampal correlations in responders, whereas non-responders showed distinct patterns, including altered endocannabinoid interactions, suggesting differential neurochemical adaptations to treatment.

## Discussion

Randomized application of mild stressors for 4 weeks resulted in increased despair, physical deterioration, and reduced body weight. These behavioral and somatic changes are comparable to the depressive-like phenotypic alterations previously described using similar UCMS protocols ^29,34–36^ and demonstrate the reliability of the model. Next, fluoxetine was administered daily for 7 consecutive days while continuing with the UCMS protocol. The selected dose of 10 mg/kg has previously been demonstrated to be effective in increasing serotonin, dopamine, and noradrenaline extracellular levels after acute administration ^15^ and to have an antidepressant effect after acute ^21^ and chronic administrations ^37^. In the present work, after short-term fluoxetine treatment, animals had a less pronounced depressive-like phenotype compared to vehicle-treated stressed mice. This effect was revealed by a healthier coat state and an increased grooming time, indicative of decreased signs of apathy, and an increased mobility time in the TST. These behavioral effects observed after 1 week of treatment at a dose of 10 mg/kg were comparable to those observed with higher doses and longer administration protocols ^29,34,38^, suggesting that a shorter treatment protocol is sufficient to elicit antidepressant-like effects in this animal model of depression. Previous preclinical studies demonstrated that chronic administration of fluoxetine distinguished two populations: those showing improved behavioral outcomes (responders) and those that did not experience a significant improvement (non-responders) ^34^. We, as previously reported ^34^, used total grooming time in the splash test, total fur coat score, and body weight as variables to distinguish these two populations. The resulting proportion of responders to non-responders in the present study (9 out of 39) is lower than the one found in previous literature (9 out of 15) ^34^. This discrepancy might be explained by the differential classification methodologies, the duration of the treatment, and/or the dose of fluoxetine. While our study employed short-term treatment, previous research has often used longer treatment periods, typically ranging from 3 to 6 weeks, to assess antidepressant-like effects ^29,34,38^. Working with shorter treatment durations allows us to expedite the treatment definition process, as behavioral effects may take longer to manifest ^13^ the underlying molecular changes occur more quickly. This enables us to identify the effects more rapidly.

The endocannabinoid system is implicated in the pathogenesis of depression and the mechanism of action of antidepressants ^39^, and in turn, cannabinoids can elicit antidepressant effects ^40^. The hippocampus is one of the brain regions with the highest CB1R density ^41,42^ associated with emotional and stress responses ^43^. Acute stress results in increased hippocampal 2-AG ^7^ and decreased AEA concentrations, reflecting a divergent response to stressors ^2^. Therefore, we analyzed changes promoted by chronic stress. UCMS did not affect 2-AG hippocampal content, but increased AEA compared to undisturbed mice. Previous studies have reported contradictory results. Indeed, chronic stress promoted hippocampal metabolic pathways associated with AEA degradation ^44^, although another study reported no alterations in hippocampal AEA following UCMS ^45^. The main differences between these previous studies and ours are the duration of the protocol (3 *vs*. 6 weeks) and the time gap between UCMS discontinuation and sacrifice (1 week *vs*. no time gap). Moreover, since samples were collected 24 h after the TST, it could be possible that the observed AEA increase in our study could be the result of an acute response to the TST stressor. These conflicting data suggest that, unlike the known effects of acute stress on AEA in the hippocampus ^46,47^, chronic stress effects on AEA must be further clarified. Monoaminergic dysfunction has been proposed as a crucial pathophysiological mechanism for depression ^48^. In our study, no significant alterations were observed as a result of chronic stress-induced depression, as previously reported ^35,36^. Unaltered serotonin levels might be the result of acute stress due to TST exposure on serotonin release ^49^, but also of the decreased expression of tyrosine hydroxylase and tryptophan hydroxylase enzymes ^36^.

In the hippocampus of control animals, endocannabinoid concentrations were positively correlated with the levels of most neurotransmitters. In contrast, exposure to UCMS disrupted these relationships, leading to an imbalance between inhibitory and excitatory neurotransmitters. These findings suggest that the modulatory role of the endocannabinoid system is impaired in chronic stress, consistent with previous reports ^2^, and might contribute to the dysregulation of other neurotransmitter systems. In addition, the endocannabinoid system is involved in synaptic plasticity processes ^50^, and chronic stress is known to alter synaptic strength and connectivity within the hippocampus ^43^. Supporting this view, previous literature linked preservation of proteins involved in vesicular functioning to stress-coping strategies ^51^. Taken together, dysfunction of the endocannabinoid system would impact hippocampal neurotransmission and synaptic plasticity, ultimately contributing to stress-related alterations.

Short-term fluoxetine significantly increased hippocampal 2-AG, mainly in mice classified as responders, suggesting that high 2-AG levels are associated with better short-term response. Excitatory neurotransmission could be involved in these changes since rapid fluoxetine-induced serotonin increase evokes endocannabinoid release through 5-HT2R activation ^52^, and endocannabinoids can act retrogradely to suppress glutamate release ^53^. In addition, chronic fluoxetine treatment results in an increase in CB1R binding capacity ^54^, strengthening endocannabinoid signaling. These findings suggest that fluoxetine effects may be partly mediated by endocannabinoid modulation, particularly through 2-AG, which influences excitatory neurotransmission. As expected, monoamine concentrations in the hippocampus and the prefrontal cortex were affected by fluoxetine. Serotonin decreased, in line with previous results ^15,35^, an effect likely mediated by the early inhibition of the 5-HT1A autoreceptors on serotonin synthesis ^22,23^. Fluoxetine also led to a comparable decrease in serotonin turnover, indicative of reduced MAO activity ^55,56^, supporting the notion that fluoxetine not only inhibits serotonin reuptake but also modulates the enzymatic pathways. There is consensus that GABA and glutamate are decreased in the cerebrospinal fluid ^57,58^ and brain ^59^ of patients and preclinical models ^44,60^ of MDD. The prefrontal cortex is highly sensitive to stress ^61^, which receives enhanced input from subcortical areas whose neuronal activity is impaired during chronic stress ^62,63^, resulting in GABAergic and glutamatergic imbalance and impairment of affective behavior ^64^. The present results show that GABA and glutamate are decreased in the prefrontal cortex of responders, suggesting their association with a better behavioral response. This effect could be mediated by the alteration of vesicle transport dynamics ^65^. Nonetheless, fluoxetine can also stimulate GABA and glutamate neurotransmission by serotonin activation of 5-HT2AR on GABAergic interneurons and glutamatergic neurons ^66^, suggesting that fluoxetine could have multiple effects.

Therefore, the depression-like phenotype induced by UCMS was associated with increased hippocampal AEA. Behavioral improvement with short-term fluoxetine was accompanied by increased hippocampal 2-AG, associated with decreased GABA and glutamate in the prefrontal cortex.

## Data Availability

All data are available in the main text or supplementary materials. Correspondence and requests for materials should be addressed to Beltrán Álvarez-Pérez, Eleni Tzavara or Rafael Maldonado.

## Acknowledgments

We thank Raquel Martín for her extensive technical assistance throughout the project, and Francisco Porrón for his expert support in the animal experiments. We also thank Rafael de la Torre, Oscar Pozo and Élida Alechaga for their expert assistance with the metabolomic studies. We are also grateful to Araceli Bergadà-Martínez, Tamara Pezzotta, Maria Ruiz Suárez, Alba Piñeiro Justo, and the Neuropharmacology laboratory for their valuable discussions and technical help.

## Author Contributions

AR-L contributed to the experimental design, performed the behavioral experiments, conducted sample collection and processing, analyzed the data, and drafted the manuscript. ECI contributed to the study design and supervised data analysis. VG participated in the project development and experimental design. MM, PR, ET, RM, and BA-P contributed to project funding and manuscript revision. PR and BA-P conceived the study and contributed to experimental design, data analysis, and manuscript writing and revision.

